# *neu*Print: Analysis Tools for EM Connectomics

**DOI:** 10.1101/2020.01.16.909465

**Authors:** Jody Clements, Tom Dolafi, Lowell Umayam, Nicole L. Neubarth, Stuart Berg, Louis K. Scheffer, Stephen M. Plaza

## Abstract

Due to technological advances in electron microscopy (EM) and deep learning, it is now practical to reconstruct a connectome, a description of neurons and the connections between them, for significant volumes of neural tissue. The limited scope of past reconstructions meant they were primarily used by domain experts, and performance was not a serious problem. But the new reconstructions, of common laboratory creatures such as the fruit fly *Drosophila melanogaster*, upend these assumptions. These natural neural networks now contain tens of thousands of neurons and tens of millions of connections between them, with yet larger reconstructions pending, and are of interest to a large community of non-specialists. This requires new tools that are easy to use and efficiently handle large data. We introduce *neuPrint* to address these data analysis challenges. neuPrint is a database and analysis ecosystem that organizes connectome data in a manner conducive to biological discovery. In particular, we propose a data model that allows users to access the connectome at different levels of abstraction primarily through a graph database, neo4j, and its powerfully expressive query language *Cypher*. neuPrint is compatible with modern connectome reconstruction workflows, providing tools for assessing reconstruction quality, and offering both batch and incremental updates to match modern connectome reconstruction flows. Finally, we introduce a web interface and programmer API that targets a diverse user skill set. We demonstrate the effectiveness and efficiency of neuPrint through example database queries.

## 1 Introduction

High-resolution EM data reveals the morphology of individual neurons and the synapses between them. By representing the neurons as nodes and synapses as edges, the resulting connectivity graph provides scientists one tool to help understand neural mechanisms in brains. Technical hurdles in generating and reconstructing connectomes from EM data limited prior studies to either small brains like *C*. elegans [1] or smaller portions of larger brains [2], [3], [4]. Despite the relatively small size, typically 1000 or fewer neurons, compared to the 100, 000 neurons in the *Drosophila* or millions of neurons in a mouse brain, deciphering the circuits formed by these neurons is challenging. The need for effective representation of complex connectomes is increasing with much larger EM datasets available[5][6] and the introduction of new methods of speeding up connectomic reconstruction, using techniques such as automatic EM image segmentation using deep learning[7].

At its simplest, a connectome is a lookup table that enables scientists to find the inputs or outputs of a given neuron. In some model organisms like *Drosophila*, one can use this information, combined with modern genetic tools, to selectively silence or monitor specific neurons to potentially infer neural mechanisms for certain behavior[8]. However, even this look-up table application poses many analysis challenges, especially for larger datasets. If the neuron type being looked up is not well-established and annotated explicitly in the database, how does one find it? Once found, many neurons have hundreds of inputs and outputs spanning large portions of the brain. Which ones are important? To minimize the need for follow-up experiments, the once simple lookup task might require a more complicated analysis of inferring the role of neurons in this population based on their connectivity and projections. This analysis will likely involve brain regions and neurons unknown to the experimenter, or often science as a whole. These challenges of interpreting large data further intensify if one wishes to infer mechanisms directly from the data, such as by trying to find underlying patterns in the connectivity graph or examining low-level motifs such as the location distribution of synapses on a given neuron.

We introduce *neu*Print as a connectome analysis framework to address the challenges of interpreting large connectome data. At its core, neuPrint is a data model for representing connectome data that provides the following advantages:

- It represents data at different levels of detail based on natural anatomical features (brain region, neuron, and synapse level) to maximize the efficiency of queries based on the needs of the users and to enable an intuitive interface consistent with the goals of the user.
- It exploits a graph database (neo4j [9]) to facilitate working efficiently with large graph data.
- It exploits brain regions (regions of interest, or ROIs) to allow users to take a top-down strategy for understanding complex data. It does this by decomposing connectome data by ROI when relevant.
- It facilitates relatively simple and straightforward queries by leveraging the expressive Cypher graph query language.
- It enables metadata properties to be flexibly added to neurons and synapses.

This data model is implemented within neo4j. We implement a connectomics interface over neo4j allowing users to access the data either programmatically or interactively through a web interface. Our web interface combines 3D visualization and a flexible plugin system to enable the rapid creation of new analysis tools to meet the demands of new usage patterns for this emerging field. Furthermore, the neuprint ecosystem can leverage other storage solutions, e.g., [10], for non-graph connectome-relevant data, such as morphological skeletons, useful in tasks such as delay modeling.

Other tools exist for analyzing connectomes [11, 12, 13, 14] [13], which between them provide an impressive collection of analysis tools. neuPrint differs by using an off-the-shelf database solution, neo4j, and its well-documented and reasonably intuitive query language, Cypher, as the medium instead of a more custom query language. Also, neuPrint emphasizes a data model that maximizes performance and interpretability with graph data in mind without as much emphasis on lower-level data and exploration. Unlike [11, 12, 14], neuPrint is not an editing tool, which again allows us to focus our design goals. An example is the use of a native graph database which has performance advantages for path queries when compared to a relational database used in other tools[12]. neuPrint does not subsume all the analysis capabilities of these editing tools. Rather, neuPrint is compatible with any workflow that leverages segmentation during editing (e.g., [2]), so that neuPrint can be used in parallel with reconstruction. This is especially important since automatic image segmentation and connectome datasets are imperfect and constantly under revision. To this end, neuPrint can help assess dataset quality and navigate reconstruction uncertainty or incompleteness.

In this paper, we first discuss our overall framework and the main data model. Then, we describe the programmer API and web application. Finally, we discuss the practical details of deploying neuPrint, and present empirical justification for our design decisions, as well as explore several example queries on a large dataset.

## 2 Storing and representing analysis data

Figure 1 shows an overview of the neuPrint ecosystem. In this section, we emphasize the representation and storage of connectome data. The next section will discuss the higher level interfaces.

**Figure 1:**
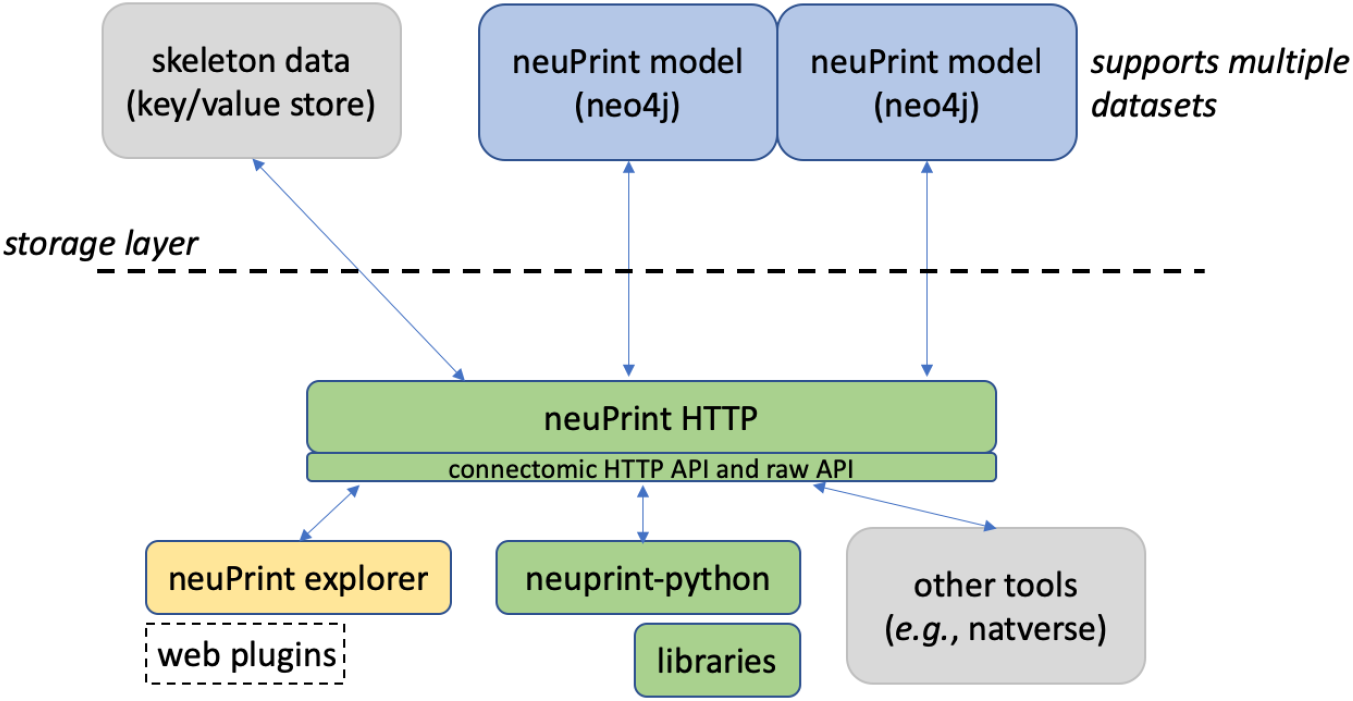
neuPrint ecosystem. The ecosystem is broadly divided into a lower-level data representation and storage above the dashed line and a higher-level interface below the dashed line.

We consider the storage of the connectomic graph and associated metadata within a graph database neo4j [9]. Presumably, other graph databases that support the graph query language *Cypher* could be compatible with neuPrint, though this has not been tested. In a graph database, nodes can access related nodes through linked lists. This is in contrast to more traditional table-based relational (SQL) databases, where finding whether a node is related to another node requires first joining those two tables together. Therefore, queries that require relationship lookups, such as path searches, are potentially much faster in a graph database.

Graph databases are often advantageous when a problem can be clearly formulated as a graph, such as the connectome. In this case a data model, or strategy for organizing the data, can be more intuitively designed for a graph database. Conversely, in a relational database, a simple graph model showing neuron nodes connected by synapse edges would require several different tables. For instance, one could have a neuron table, an synapse or edge table, a neuron property table, and an edge property table. Graph databases and other so-called NoSQL databases tend to not require an exact schema, meaning that it is easy to add new relationship types on pre-existing data models. This is advantageous in connectomics as we anticipate the need to adapt quickly to new analysis requirements.

The EM connectomic dataset involves other data useful for analysis that are not ideally suited for a graph database. For larger storage objects, like a neuronal skeleton (which is a simplified ball and stick representation of a neuron’s morphology) and ROI surface meshes, we leverage simpler key/value stores where one retrieves a value by using a specific key or address. While one can reasonably store a series of skeleton nodes in a graph database, we found that most analyses involving skeletons required the whole skeleton meaning that a simple fetch of the whole data structure was sufficient, and more time and space efficient.

In the following, we present the neuPrint graph data model and then explain the ramifications.

### 2.1 Data model

We illustrate how the data is organized in the graph database in Figure 2. There are five major node types or labels denoted by the syntax “:”. In neo4j these labels help partition the nodes into different groups. `:Neuron` and `:Synapse` nodes are two obvious aspects of a connectome. Neurons contain several properties (with more details in the Appendix). The bodyId is a mandatory field and is a unique identifier for the given neuron. Other fields are required as indicated in the figure. neo4j allows one to index different properties for a node label, reducing querying time at the cost of more disk storage. Synapses contain x,y,z location, which are indexed properties that can be accessed using neo4j’s spatial querying capabilities. The synapses for a given neuron are grouped under different nodes called `:SynapseSet`. There is a synapse set that groups all the synapses for each connection for each neuron. The `:Meta` node type provides top-level information about the database.

**Figure 2:**
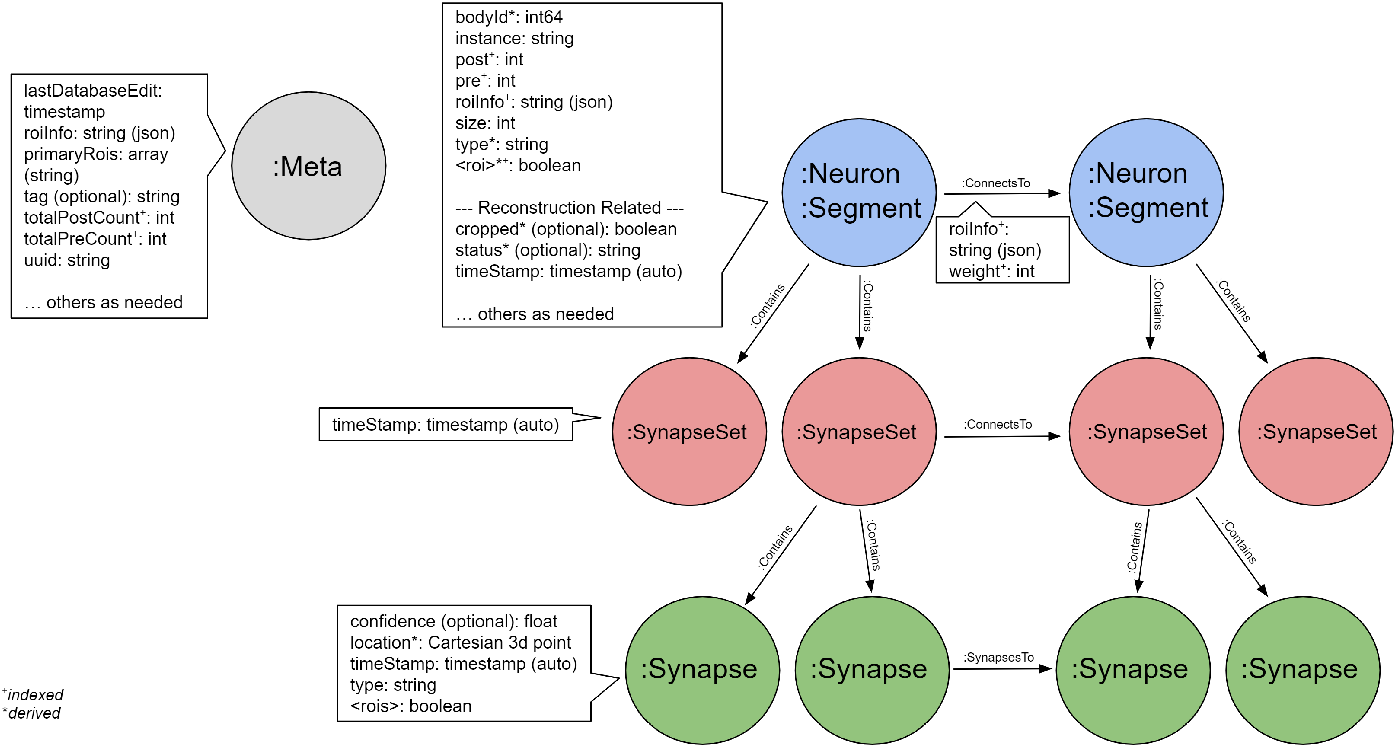
neuPrint graph data model. This shows the various node types and properties used for storing data relevant for connectome analysis.

The data model also has a `:Segment` node label. As noted earlier, neuPrint was designed to be useful for quality control for connectome datasets. The current best practice for creating a connectome requires using automatic image segmentation. This typically creates, in addition to large neuronal pieces, many small segment fragments that could still be edited and joined to a neuron. A `:Segment` node is superset of `:Neuron`. The criteria for labeling a `:Segment` as also a `Neuron` depends on the usage patterns of the database and what is considered to be relevant for most queries.

Between these different node types, we define several relationship types. Prominently, the segments (or neurons) and synapse sets are connected via a `:ConnectsTo` property. Individual synapses are linked together through a `:SynapsesTo` property. We use the `:Contains` relationship to define the synapse sets that each segment contains and the synapses that each synapse set contains.

Region information is encoded in the data model at multiple levels. Each synapse has a boolean value for each ROI it resides in. Since ROIs are hierarchically defined, several such values may be set. The synaptic ROI information is also aggregated over segments and connections and is stored in the `roiInfo` field. This enables users to easily extract the number of synapses per region for a segment or connection.

For each node label, we also partition the node using a dataset-specific indicator. For instance, a neuron for the dataset named “x”, would be “:x_Neuron”. In this manner, we can support multiple datasets in the same database. Queries can be made across datasets or targeted to a specific dataset.

### 2.2 Design considerations

This section explains the motivation for some of the data model design decisions.

The primary goal of the data model design was to encourage top-down use of the data model and to allow users to exploit region information extensively. The most common queries will only involve neuron connections, which exists as a redundant higher-level representation in our model. An alternative data model design could require the user to extract neuronal connectivity by traversing every synapse between two neurons – a slower, and more complicated query. The ROI information is similarly encoded at multiple levels to facilitate query performance and ease of use. Even though it is possible to compute region statistics from the synapse points, it is faster and easier to find neurons in certain regions and get basic region statistics by simply querying information available at the neuron and neuron connection level.

Creating a connectome from automatic segmentation often requires merging several smaller segments together into large segments [15]. Based on prior experience, a segmentation contains many more segments than neurons. This observation motivated creating `:Segment` and `:Neuron` labels as a mechanism of partitioning the most important segments. By restricting most analyses to `:Neuron` labels, queries that require linear scans can operate much faster. We also have an optional property status tracking the quality of a given neuron reconstruction. Finally, each synapse contains a confidence field, typically computed by automatic synapse prediction, that can be used to model confidence for certain neuron connections.

#### Future considerations

The proposed data model can be extended in many different ways. By allowing multiple datasets in the same neo4j (by using the dataset prefix for each node label), one could add specific relationships between related neurons across datasets. Also, if there are many more property types required for a segment, it might make sense to create a separate `:SegmentProperty` node. We could also extend the model to accommodate other cell ultra-structure. For instance, we would add a _:MitoSet_ to link to `:Mitochondrion` nodes for each neuron. Finally, a relationship type like `:Merge` could indicate segments that could be grouped together.

The current strategy for embedding `roiInfo` at the connection level and segment level is convenient but clumsy. Because neo4j does not support map datatypes (where a list of keys can have an associated value), the data is encoded as a JSON string. This data cannot be indexed in a meaningful way and requires decoding the JSON when used as a filter within a query. One could encode region breakdown per ROI with the introduction of explicit `:Region` nodes. While this might be more idiomatic, it leads to more complex user queries, hence our current design decision. Finally, the current data model treats each ROI or brain region separately. If the ROIs available form a hierarchy, one could presumably simplify roiInfo by providing stats only for the ROIs at the lowest level of the hierarchy.

## 3 Interfacing with neuPrint

To enable unified access to the underlying data model and other connectomic data, such as neuron skeletons, we provide a software layer, neuPrintHTTP. neuPrintHTTP provides a mostly read-only connectomic-specific interface that allows users to make HTTP requests that then call the underlying neo4j database or other storage engines. It also simplifies querying within a given dataset. As previously noted, each node label actually encodes the dataset name, such as <`dataset`>_<`node label`>. With neuPrintHTTP, the user can direct queries to a given dataset without having to provide dataset-specific labels.

neuPrintHTTP is designed in the language *Go* to exploit convenient concurrency semantics, so it can handle several parallel requests efficiently. Furthermore, the backend of the software layer abstracts the storage into different technology-specific plugins. For the non-graph data, plugins exist to access DVID [10] and a generic key-value database. Other databases that can satisfy the interface requirements can be easily added, such as Google storage or Amazon S3. neuPrintHTTP also supports authentication with Google OAuth and provides options to make the data read only for anyone, or to restrict access to a set of authorized users. neuPrintHTTP also has a mode to enable database writes for given admin-level authorized users.

### 3.1 neuPrint Web Explorer

In many cases, users might prefer an interactive visual interface with neuPrint over the use of APIs. To this end, we introduce neuPrintExplorer. neuPrintExplorer is a web application that interacts with neuPrintHTTP written using the modular web framework called REACT. It provides a series of different common analysis queries within different plugins. Each plugin is a gateway into accessing the data. At its simplest, most queries involve displaying some table of information based on a simple database request to neuPrintHTTP. In addition, many of these plugins create visuals such as charts that breakdown neuron or connections (see Figure 3) to separate brain regions or provide links to access other parts of the dataset.

**Figure 3:**
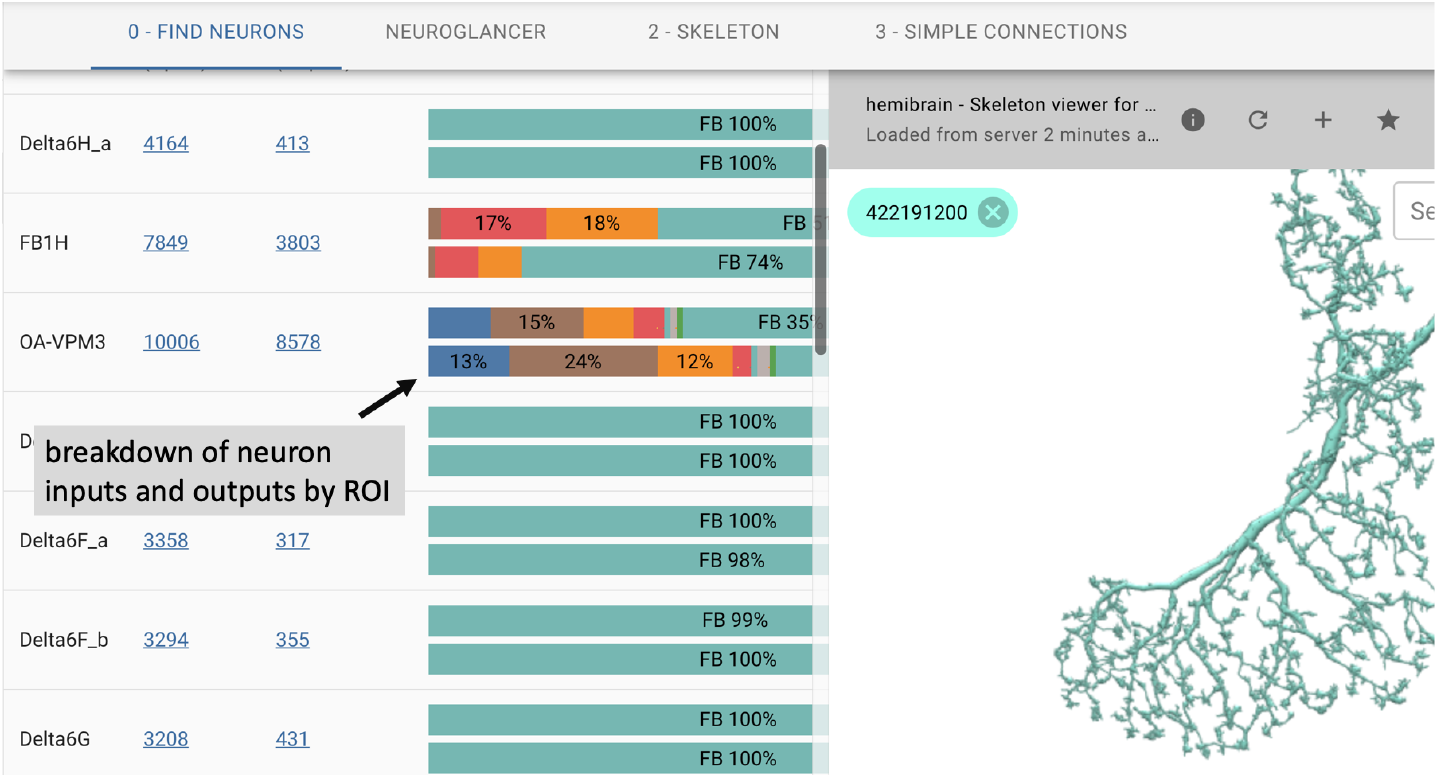
neuPrintExplorer web application. Queries generate tables of results. Visualizations exist to see 3D neurons and to help break down the complexity of the data.

As shown in Figure 3, the web application supports 3D visualization of neurons by embedding the skeleton viewing tool called SharkViewer (*https://github.com/JaneliaSciComp/SharkViewer*). This allows users to see the morphology (shape) of given neurons (fetching the data from the neuprintHTTP’s skeleton endpoint) and also the arrangement of synapses on these neurons. We have also implemented a REACT wrapper around the powerful web application neuroglancer (*https://github.com/google/neuroglancer*), designed for browsing EM datasets. This means that we can embed neuroglancer within our application and enable users to find neurons in neuroglancer based on interactions in neuPrintExplorer. While neuPrint is designed for analysis only, part of the connectome reconstruction process sometimes requires users to add annotations or comments on the underlying dataset. By supporting neuroglancer, neuPrintExplorer provides a gateway for lower-level exploration and annotation if needed.

Architecturally, neuprintExplorer is a single-page application written using REACT/Redux which allows us to leverage other open source components such as d3 for graphics or material-ui for the UI. We also designed the system to be modular by providing a plugin system that allows new queries and views to be added without modifying the core code. There are example plugins and instructions on how to create a new plugin at *https://github.com/connectome-neuprint/neuPrintExplorerPlugins*.

#### Example plugins

Some of the plugins we have created for the most common tasks are:

- Simple connections: find inputs or outputs for a neuron ordered by connection strength. This plugin implements the simple lookup table.
- Find neurons: find neurons in the dataset by name and/or by the regions that they have inputs and outputs.
- Shortest paths: find all shortest directed paths from one neuron to another and display the local connectivity graph. This query is generally very efficient except for very deep (or non-existent) paths. A timeout is set for a few seconds.
- Find Similar Neurons: find neurons whose inputs and outputs intersect ROIs similar to the provided neuron.
- Cell type: show all neurons of the same cell type to evaluate the connection similarity between neuron of the same type (this is an example of a more complicated query compared to a simple Cypher request).
- Brain region connectivity: show how the brain regions connect to each other by considering the neurons that go from one region to another.
- Common connectivity: view inputs and outputs common to a set of neurons.
- Custom: allow users to execute custom Cypher queries
- Partner completeness (reconstruction QC tools): examine how fragmented the inputs or outputs are for a neuron.
- Completeness (reconstruction QC tools): show the percentage of segments for each brain region that are traced neurons.

To facilitate learning Cypher, relevant plugins provide information on the specific Cypher query made.

### 3.2 Programmer APIs

As mentioned, neuPrint provides an HTTP, or REST, interface to enable programmatic access to the underlying data. Given the diversity of analysis requirements, many of which are currently unknown, we have aimed for a lean HTTP API from which more specific capabilities can be written, such as in our python library or the R packages in natverse [16].

The most basic API endpoint provides direct query access to the neo4j interface through the Cypher query language. Cypher shares semantic similarities with SQL and is intended to provide a mostly intuitive language to query a graph database. Below is an example of a query that returns all downstream partners, m, from a given neuron, n with body id 123, with more than 10 connections.
**MATCH** (n :Neuron)-[x :Connections]->(m)
**WHERE** n.bodyId=123 AND x.weight >10
**RETURN** m.bodyId

Most Cypher queries have these three components. A **MATCH** statement identifies the pattern to be found. In this case, that is *n* with a connection to *m* (the direction of the connection is indicated by the arrow). A **WHERE** statement applies filters to the above MATCH statement. Here we restrict the match to a neuron *n* with unique body id 123 and with connection weight or strength greater than 10. Finally, the **RETURN** statment provides the results back to the user, which in this case is just the partner(s) body id. There are several online resources for learning Cypher. We will show a few other examples later in the results section.

In addition to this Cypher interface, neuprintHTTP subdivides its HTTP API into different categories. For example, there is a sub-category called “dbmeta” for database meta information and one called “npexplorer” to provide convenient wrappers for common connectome queries used in the web interface defined below, such as finding neurons that intersect certain regions. This connectomics interface is a work in progress. We plan to extend the interface to provide a simplified wrapper around the most common types of Cypher queries, as access patterns are better understood. More information on this interface can be found at *https://github.com/connectome-neuprint/neuPrintHTTP*. More information on the python API can be found at *https://github.com/connectome-neuprint/neuprint-python*.

## 4 Initializing, updating, deploying neuPrint

To use the neuPrint ecosystem requires loading data into neo4j and configuring the relevant backend storage solutions to be accessible to neuPrintHTTP. In this section, we focus on how to load data into the graph database and how to manage connectome datasets that are being actively revised and reconstructed. In particular, we hightlight the light-weight data requirements and ability for neuPrint to be updated via a light-weight incremental interface.

### 4.1 Ingesting data into the neuPrint graph data model

The current ingestion system (documented in more detail at *https://github.com/connectome-neuprint/neuPrint*) involves initializing neo4j with a series of CSV files. The files are formatted to minimize computation in neo4j to speedup ingestion, moving the computational burden to creating these CSV files. The motivation here is that neo4j is typically deployed on a single server, while these CSV files can be generated outside this environment with a multi-process initialization approach. Creating this initialization routine to enable efficient processing of very large datasets is a work in progress and currently only supports having one dataset per database (even though our data model and interfaces allow datasets to share the same database).

At a high-level the ingested data is relatively compact compared to the size of the underlying EM dataset. It involves the following components.
- A list of synapse points where each point indicates the parent body id and ROI(s).
- A edge list that shows how the synapse points are connected.
- A neuron or segment id list that contains associated metadata for each body id.
- Other metadata such as the brain regions or ROIs available and the version of the dataset.

### 4.2 Incrementally updating the neuPrint graph data model

The previous sub-section detailed the initial ingestion process which is streamlined to enable fast, one-time creation of a neo4j instance. As previously mentioned, the neuPrint ecosystem is designed to be compatible with modern connectome reconstruction workflows that use image segmentation. To this end, neuPrint is mostly decoupled from reconstruction workflows except for an incremental interface for updating the underlying data model.

Figure 4 shows the architecture for incrementally updating neuPrint. The key feature is that we require access to only the changes to the dataset, such as segment merge and split events, published to a centralized log manager, which in our case is Apache Kafka. We have light-weight services written in Python that listen for changes recorded to this log and modify the neuPrint data model using targeted Cypher statements. For example, a user can modify segmentation data using a tool like [14], which modifies data managed by DVID [10]. DVID then emits log messages to Kafka, which our Python services then consumes and updates neuPrint graph data through neuPrintHTTP.

**Figure 4:**
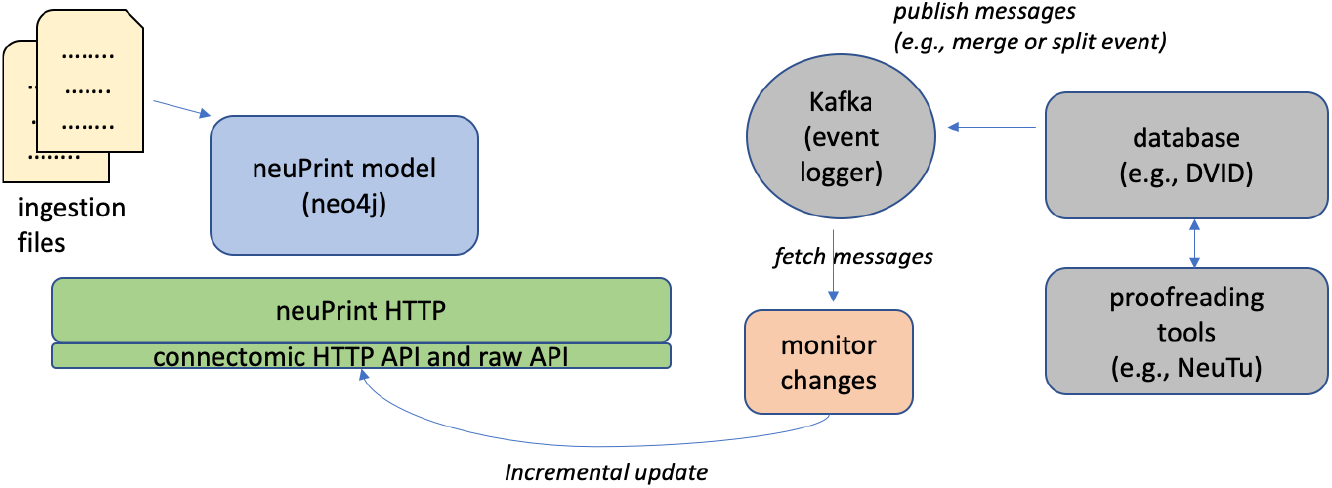
Initializing and updating the neuPrint graph data. The database is initialized by ingesting data via a series of CSV files. To update the data incrementally, a service monitors dataset changes recorded in Apache Kafka and makes incremental updates through neuPrintHTTP.

### 4.3 Deploying and managing multiple datasets

Graph databases perform best when the graph fits into one server’s shared memory. This is because queries can easily span disparate parts of the graph, given the so-called small world properties exhibited by many graphs. Fortunately, the storage requirements for our data model is quite compact compared to the original EM data and is strongly related to the number of synapses in the dataset. Extrapolating from results provided in the next section, one could expect 100s of millions of connections would be manageable on a single large server. Even larger datasets could presumably be served using fast backup SSDs with some performance penalty. A more extreme solution to even larger datasets could involve splitting synapse annotations out of the graph data model.

As previously mentioned, neuPrint allows multiple datasets to be stored in one graph database. We also allow datasets to be stored across distinct neo4j databases, since in most cases the queries we are interested typically involve examining one database at a time. This is helpful for performance reasons to ensure more dedicated memory for each dataset. We use this capability to support checkpoint management. If a dataset is actively being reconstructed, snapshots can be created for each version of the dataset and these versions are stored in different neo4j instances and orchestrated by neuPrintHTTP.

## 5 Results

In this section, we provide some insights on the performance characteristics of our system. Comparing neo4j with other relational databases is beyond the scope of this paper. Rather, we try to first demonstrate the effectiveness of our data model and then show that common queries achieve interactive performance (*i.e*. queries are under a few seconds). The example queries explored also serve as documentation for different use cases.

We make available two neuPrint datasets at *https://neuprint-examples.janelia.org*: the fly medulla seven-column dataset [2] and fly mushroom body dataset [3]. Storage and ingestion performance characteristics are provided in Figure 5 for those datasets and a larger unpublished dataset with 10s of millions of synaptic elements. The ingestion was performed on a machine with 256GB of memory and 20 processors.

**Figure 5:**
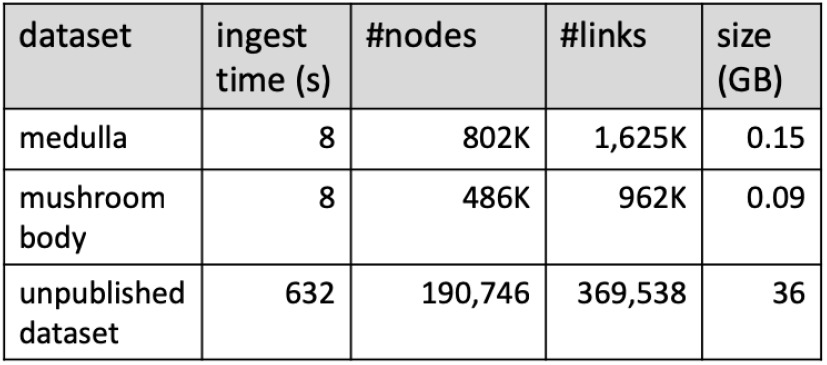
neuPrint graph representation performance. The ingestion and storage requirements for three example datasets.

The smaller two datasets, which were some of the largest connectomes produced when they were published, load in only a few seconds. Notably, the much larger dataset’s load time shows that performance scales well with increased size. As mentioned earlier, we format CSV files to streamline ingestion into neo4j. Notably, the CSV files are about the same size as the neo4j database on disk. The relatively small database sizes for even the larger dataset suggest that a much larger dataset could reside completely within memory on a large server. Of course, working with larger graphs that do not fit in memory is possible by leveraging SSD storage with some consequences for performance.

For the next two subsections, we evaluate runtime performance of various queries on the much larger, unpublished dataset. The graph data was stored in a cloud VM with memory capacity large enough to hold the entire dataset. Given the remote location of the server, each query includes several milliseconds of latency to access it. All rumtime numbers reported are a result of averaging runtimes of over 50 independent queries.

### 5.1 Performance Decisions

As discussed in Section 2.2, the graph data model was designed to facilitate ease of use and runtime efficiency. We evaluate some of the data model design decisions through three different scenarios shown in Figure 6. For each scenario, we query the database in two ways: optimized queries that leverage the full data model and less-optimized queries that assume a more simplistic data model. In all cases, we notice that the optimized queries are at least 2x faster. We provide more details in the following paragraphs. While the absolute runtime is relatively fast for two of these scenarios, programs may issue 100s of queries in a short-time and 2x runtime improvements would be more noticeable.

**Figure 6:**
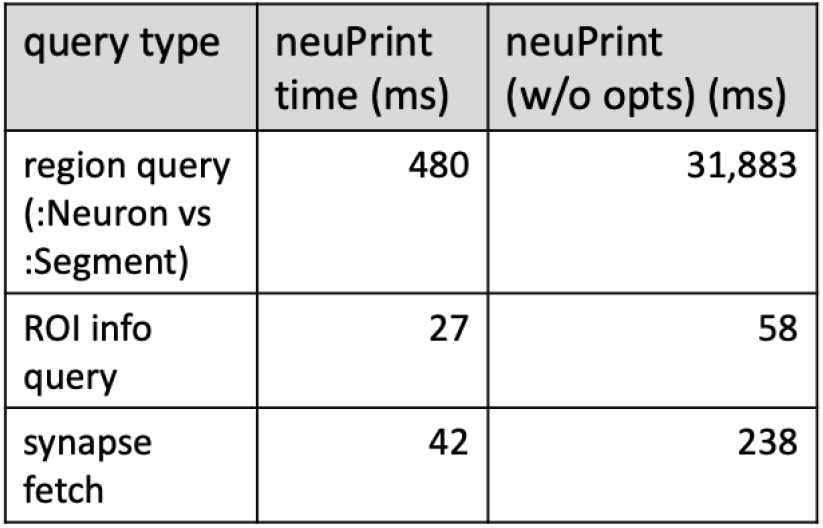
Optimized vs non-optimized query performance. Shows examples of different types of queries and the performance in milliseconds of querying using our full data model versus a simplified subset of the data model.

#### Test 1: Segment vs neuron

This test evaluates the decision to partition a subset of `:Segment` nodes into `:Neuron` nodes. The motivation was to focus queries on the more important, but less numerous `:Neuron` nodes. The below queries try to count the number of neurons or segments over a certain size in each region.

Counting neurons for an `ROI`.
**MATCH** (n :Neuron)
**WHERE** n.`ROI` AND n.size > 100000000
**RETURN** count(n)

Counting segments for an `ROI`.
**MATCH** (n :Segment)
**WHERE** n.`ROI` AND n.size > 100000000
**RETURN** count(n)

In this example, the large performance disparity is also due to ROI names being indexed to `:Neuron`. But even if we force a linear scan through all `:Neuron` nodes (which involves 1/100 the number of total segments), we still observe queries under one second. We could in principle create indices for every property for a segment, but each index comes with a storage cost which is magnified because there are many more segments than actual neurons. Therefore having a special `:Neuron` designation potentially reduces the database size and can improve performance.

#### Test 2: roiInfo

This test checks the performance of using the `:ConnectsTo` property, `roiInfo`, versus examining the ROI information by inspecting individual synapses. The following two queries examine a given neuron connection to see if the connection is in a given ROI. The first uses the `roiInfo` property on the connection edge. The second one inspects region information by looking at all the synapses within a synapse set.

Checks if connections exist between `body1` and `body2` in a certain `ROI`.
**MATCH**(n :Neuron {bodyId: `body1`})-[x :ConnectsTo]->(m :Neuron {bodyId: `body2`})
**RETURN** EXISTS(apoc.convert.fromJsonMap(x.roiInfo)[“`ROI`”])

Checks if connections exist `body1` and `body2` in a certain `ROI` without using denormalized roi information.
**MATCH** (n :Neuron {bodyId: `body1`})-[:Contains]->(:SynapseSet)-[:ConnectsTo]->(ss :SynapseSet)<-[:Contains]-(m :Neuron {bodyId: `body2`})
**MATCH** (ss)-[:Contains]->(syn :Synapse)
**WHERE** syn.`ROI`
**RETURN** true LIMIT 1

While using the `roiInfo` property results in a 2x faster query, more importantly, the first query is much more compact and easier to understand.

#### Test 3: Accessing synapses through synapse sets

This test motivates synapeset sets (`:SynapseSet`) as a mechanism of grouping synapses together. In general, by grouping synapses together, we can minimize the number of edges on a given :Segment in the graph model, presumably accelerating queries involving segments. In our model, `:SynapseSet` nodes are specific to each connection between two segments. The queries below provides an example of downloading all synapses for a given connection either using synapse sets or by determining the relationships by exploring the lowest level `:SynapsesTo` relationship.

Retrieving post-synaptic sites for connections with `:SynapseSet`.
**MATCH** (n :Neuron {bodyId: `body1`})-[:Contains]->(:SynapseSet)-[:ConnectsTo]->(ss :SynapseSet)<-[:Contains]-(m :Neuron {bodyId: `body2`})
**MATCH** (ss)-[:Contains]->(syn :Synapse)
**RETURN** syn.location, syn.confidence

Retrieving post-synaptic sites for conneections without `:SynapseSet`.
**MATCH** (n :Neuron {bodyId: `body1`})-[:Contains]->(:SynapseSet)-[:Contains]->(:Synapse)-[:SynapsesTo]->(syn :Synapse)<-[:Contains]-(ss :SynapseSet)<-[:Contains]-(m :Neuron {bodyId: `body2`})
**RETURN** syn.location, syn.confidence

Both Cypher queries are slightly more complex to express. However, the synapse sets enable much faster performance.

### 5.2 Example queries

In this section, we survey different analysis use cases and provide a sense of runtime performance averaged over several runs. Notably, most queries require only a fraction of a second. The most complex query involves looking for all 3-hop paths for several random pairs of neurons. The average runtime for this query is under 5 seconds but we note wide variance with many queries finishing under a second and some taking around 30 seconds.

Example 1: sum connection weight of partners that are traced.
**MATCH** (n :Neuron {bodyId: `bodyid`})-[x :ConnectsTo]->(m :Neuron)
**WHERE** m.status=“Traced”
**RETURN** sum(x.weight)

**Figure 7:**
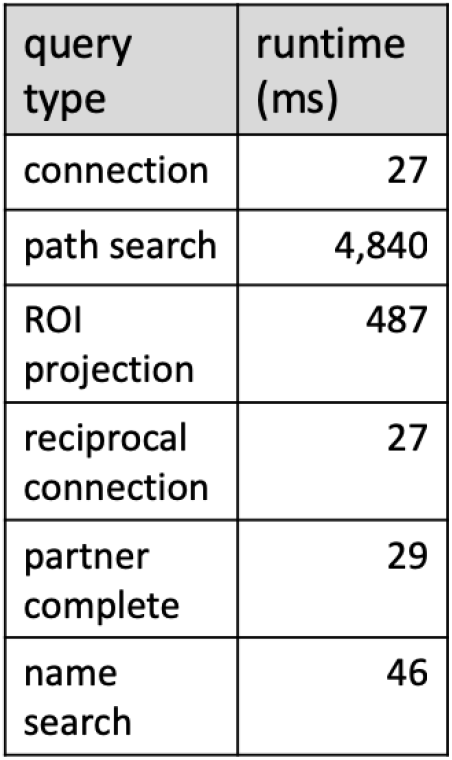
Performance of example queries. Runtime in milliseconds is averaged over several different queries of the same type.

Example 2: find all paths up to 3 in length between two neurons with >=5 connections.
**MATCH** p =(n :Neuron {bodyId: `body1`})-[x :ConnectsTo*..3]->(m :Neuron bodyId: *body2*
**WHERE ALL** (x in relationships(p) WHERE x.weight >= 5)
**RETURN** count(p)

Example 3: find neurons projecting from one region to another.
**MATCH** (n :Neuron)
**WHERE** n.`ROI1` AND n.`ROI2`
**WITH** apoc.convert.fromJsonMap(n.roiInfo) AS info, n
**WHERE** info[`ROI1`].post >0 AND info[`ROI2`].pre >0
**RETURN** count(n)

Example 4: Find reciprocal connections.
**MATCH** (n :Neuron {bodyId: `body1`})-[x :ConnectsTo]->(m :Neuron {bodyId: `body2`})
**WHERE** (m)-[:ConnectsTo]->(n)
**RETURN** true

Example 5: Find reconstruction incompletness for a neuron’s outputs (returns the distribution of reconstruction statuses including which connections are “Traced”).
**MATCH** (n :Neuron {bodyId: *bodyid*})-[x :ConnectsTo]->(m :Segment)
**RETURN** m.status as status, sum(x.weight) AS total

Example 6: Lookup neuron type with regular expressions.
**MATCH** (n :Neuron)
**WHERE** n.type=~“*prefix*.*”
**RETURN** count(n)

## 6 Conclusions

We introduce the neuPrint ecosystem in this paper as a mechanism to aid in large-scale analysis of EM connectomes. The central component of neuPrint is the graph data model that stores the data in an efficient manner, accessible to a variety of users and use cases. To this end, we highlight both a custom interactive web interface and programmer interfaces. Our results show that our database enables a diverse set of queries with a dataset containing millions of synaptic connections.

Creating platforms and resources for large EM connectomic datasets pose different challenges than other neuroscience resources, such as VirtualFlyBrain[17], the Allen Brain Map[18], or the Mouse Light project[19]. These resources typically involve the collection of several (often smaller) datasets that are combined to form canonical atlases. In the case of connectomes, a single dataset is expensive to acquire and often very large. The notion of a canonical connectome atlas is less meaningful currently. As such, neuPrint emphasizes access to specific datasets rather than a general compilation of many datasets.

An EM image volume often contains much more information than simply neurons and synapses. Future work will involve incorporating information about the location and arrangement of various sub-cellular organelles into the data model. We believe that tools like neuPrint will be critical for managing the complexity of such rich datasets, especially as the means for extracting this information automatically become more reliable.

## Acknowledgements

We would like to thank the entire Janelia FlyEM team, many who provided extensive feedback and support for neuPrint. We thank Emily Joyce for help with documentation, Eric Trautman and Rob Svirskas for software support, and Reed George for manuscript feedback.

## 7 Appendix

This section defines many of the various properties shown in Figure 2.

**:Segment (:Neuron) nodes**

- bodyId: a unique number for each distinct segment
- pre: Number of pre-synaptic sites on the segment
- post: Number of post-synaptic sites on the segment
- type: Cell type name for given neuron (if provided)
- instance: String identifier for a neuron (if provided)
- size: Number of voxels in the body
- roiInfo: JSON string showing the `pre` and `post` breakdown for each ROI the neuron intersects.
- <roi>: This property only exists for the ROIs that intersect this segment
- status: Reconstruction status for a neuron. By convention, we broadly consider proofread neurons as being “Traced”.
- cropped: Since datasets often involve a portion of a larger brain, cropped indicates that a significant portion of a neuron is cut-off by the dataset extents. By convention, all “Traced” neurons should be explicitly noted whether they are cropped or not.

**:Synapse nodes**

- confidence: floating point value depicting confidence in synapse annotation (higher score means more confidence).
- location: x,y,z location for the synapse. This location will be on its parent segment. The location is indexed to enable fast spatial queries.
- <roi>: This property only exists for the ROIs that intersect this synapse

**:ConnectsTo relationship (between two :Segment nodes)**

- weight: number of synapses (or weight) between the two neurons
- roiInfo: JSON string showing the `pre` and `post` breakdown for each ROI the connection intersects

**:Meta node**

- uuid: some version for this dataset. This could be a DOI. Similar to a GIT commit ID used in software development.
- tag: a release tag similar to the tags provided for software releases (e.g., “v1.0”)
- primaryRois: an array of ROIs that make up the primary ROIs (or default level of ROIs in the ROI hierarchy). This is useful for various plugins in neuPrint explorer.
- roiInfo: JSON string showing the `pre` and `post` breakdown for each ROI.

